# *In vivo* analyses reveal rapid and permissive lipid transport between the ER and mitochondria

**DOI:** 10.64898/2026.06.18.733118

**Authors:** Paul Montmayeul, Samantha Voguin, Catherine Albrieux, Hanna Kulyk, Lindsay Peyriga, Floriant Bellvert, Lucas Place, Marion Schilling, Juliette Jouhet, Alexandre Toulmay, William A. Prinz, Morgane Michaud

**Author notes:** Equal contribution (Paul Montmayeul, Samantha Voguin).

## Abstract

Interorganelle lipid transport is essential for mitochondrial membrane biogenesis and function, yet its kinetics and substrate selectivity remain poorly understood *in vivo*. Here, we developed two complementary approaches to quantify lipid trafficking from the endoplasmic reticulum (ER) to mitochondria in yeast. Metabolic labeling combined with organelle fractionation revealed that newly synthesized phospholipids rapidly accumulate in mitochondria, with 20-35% of newly synthesized molecules detected in mitochondrial fractions within minutes of synthesis. To directly quantify lipid flux, we established a synthetic transport assay based on the production of heterologous galactolipids absent from yeast. This approach revealed an ER-to-mitochondria transport flux of approximately 2.6 x 10^5^ lipid molecules per cell per minute. Remarkably, galactolipids were transported with high efficiency despite their absence from fungal membranes, indicating limited substrate selectivity of ER-mitochondria lipid transport pathways. Together, these complementary assays provide quantitative tools to investigate intracellular lipid transport and reveal the rapid and permissive nature of lipid exchange between the ER and mitochondria.

**Summary:** Using complementary metabolic labeling and synthetic lipid reporter assays, we quantitatively measured ER-mitochondria lipid transport in yeast. Our results reveal rapid lipid exchange, high transport fluxes and limited substrate selectivity, indicating that mitochondrial lipid trafficking efficiently accommodates structurally diverse membrane lipids.

## Introduction

Mitochondria are essential organelles whose biogenesis and function depend on the continuous exchange of lipids with other cellular membranes. Because the vast majority of mitochondrial phospholipids are synthesized outside mitochondria, the maintenance of mitochondrial membrane composition requires efficient lipid transport from the main site of lipid synthesis, the endoplasmic reticulum (ER), and other organelles (Mavuduru et al., 2024; Tamura et al., 2020; Acoba et al., 2020). In budding yeast, mitochondria also play a central role in cellular lipid metabolism by serving as the primary site of phosphatidylethanolamine (PE) synthesis through the decarboxylation of phosphatidylserine (PS) by the phosphatidylserine decarboxylase Psd1, located in the mitochondrial inner membrane (Gaigg et al., 1995; Achleitner et al., 1995; Horvath and Daum, 2013). These observations highlight the importance of interorganelle lipid trafficking for mitochondrial and cellular homeostasis.

Lipid transport to mitochondria occurs predominantly through non-vesicular mechanisms at membrane contact sites (MCSs), specialized regions where two organelles are closely apposed (Mavuduru et al., 2024; Tamura et al., 2020; Acoba et al., 2020; Prinz et al., 2019). In yeast, mitochondria establish extensive contacts with both the ER and the vacuole (Gaigg et al., 1995; Mavuduru et al., 2024; Lang et al., 2015; Hönscher et al., 2014). At these sites, lipid movement is thought to be mediated by lipid transfer proteins (LTPs), which contain hydrophobic cavities capable of shielding lipids from the aqueous cytosol (Hamaï and Drin, 2024; Reinisch et al., 2025). LTPs can be broadly classified into shuttle-type proteins, which transfer individual lipid molecules between membranes, and bridge-type proteins, which form hydrophobic channels that likely support bulk lipid transport. In yeast, despite the identification of several lipid transport systems, including the ER–mitochondria encounter structure (ERMES) complex and the bridge-like transporter Vps13 (Petrungaro and Kornmann, 2019; Mavuduru et al., 2024; Kornmann et al., 2009; Lang et al., 2015), the mechanisms that govern the rate, directionality, and substrate selectivity of mitochondrial lipid transport remain poorly understood.

A major obstacle to addressing these questions is the lack of quantitative approaches for measuring lipid transport *in vivo*, particularly in yeast, despites significant progress these last years (Hamaï and Drin, 2024; Iglesias-Artola et al., 2025). Analyses of mitochondrial lipid composition have provided valuable insights into lipid homeostasis but generally cannot distinguish defects in lipid transport from alterations in lipid synthesis, remodeling, or degradation. Historically, mitochondrial PS decarboxylation by Psd1 has been used as a proxy for ER-to-mitochondria lipid transport (Achleitner et al., 1995; Kornmann et al., 2009; Lahiri et al., 2014; Hamaï and Drin, 2024). Although this approach revealed rapid conversion of newly synthesized PS into PE, its interpretation has become more complex following reports that a fraction of Psd1 can localize to the ER under specific growth conditions (Friedman et al., 2018). More recently, the METALLIC strategy enabled the monitoring of phosphatidylcholine (PC) transport through dual isotopic labeling of lipid headgroups and acyl chains (John Peter et al., 2022; John Peter and Kornmann, 2024). Nevertheless, broadly applicable methods for quantifying lipid transport kinetics and substrate selectivity *in vivo* in yeast remain scarce.

Importantly, lipid trafficking must be tightly coordinated with lipid synthesis and turnover to meet the changing demands of cell growth, membrane biogenesis, and stress adaptation. Therefore, understanding mitochondrial lipid transport requires approaches that capture these processes dynamically and within the context of whole-cell lipid metabolism. Here, we developed two complementary *in vivo* assays to analyze lipid transport from the ER to mitochondria in *Saccharomyces cerevisiae*. By combining short-term metabolic labeling with a synthetic lipid reporter system, we measured the extent, kinetics, and substrate selectivity of mitochondrial lipid trafficking and uncovered a rapid and highly permissive lipid transport network between the ER and mitochondria.

## Results and discussion

### In vivo measurement of lipid transport to mitochondria by metabolic labeling

To quantify lipid transport from the endoplasmic reticulum (ER) to mitochondria *in vivo*, we developed a metabolic-labeling assay that monitors the appearance of newly synthesized phospholipids in isolated mitochondria (**Fig. 1A**). The strategy relies on labeling ER-derived phospholipids with a common precursor and measuring the incorporation of radioactivity into total cellular lipids and mitochondrial lipids over time. Several precursors can be used to label phospholipid synthesis, including glycerol-3-phosphate, fatty acids, and acetate. We found that [^3^H]-acetate provided the most robust signal, with linear incorporation into most phospholipid classes during a 10-30 min labeling period in both whole-cell and mitochondria fractions (**Fig. S1 and S2**). To facilitate mitochondrial isolation, labeling was performed in yeast spheroplasts. Because cell wall digestion is a stressful treatment altering mitochondrial morphology, spheroplasts were allowed to recover for 1 h in osmotically stabilized medium before labeling. This regeneration step restored the normal mitochondrial network morphology (**Fig. 1A**).

**Figure 1.**
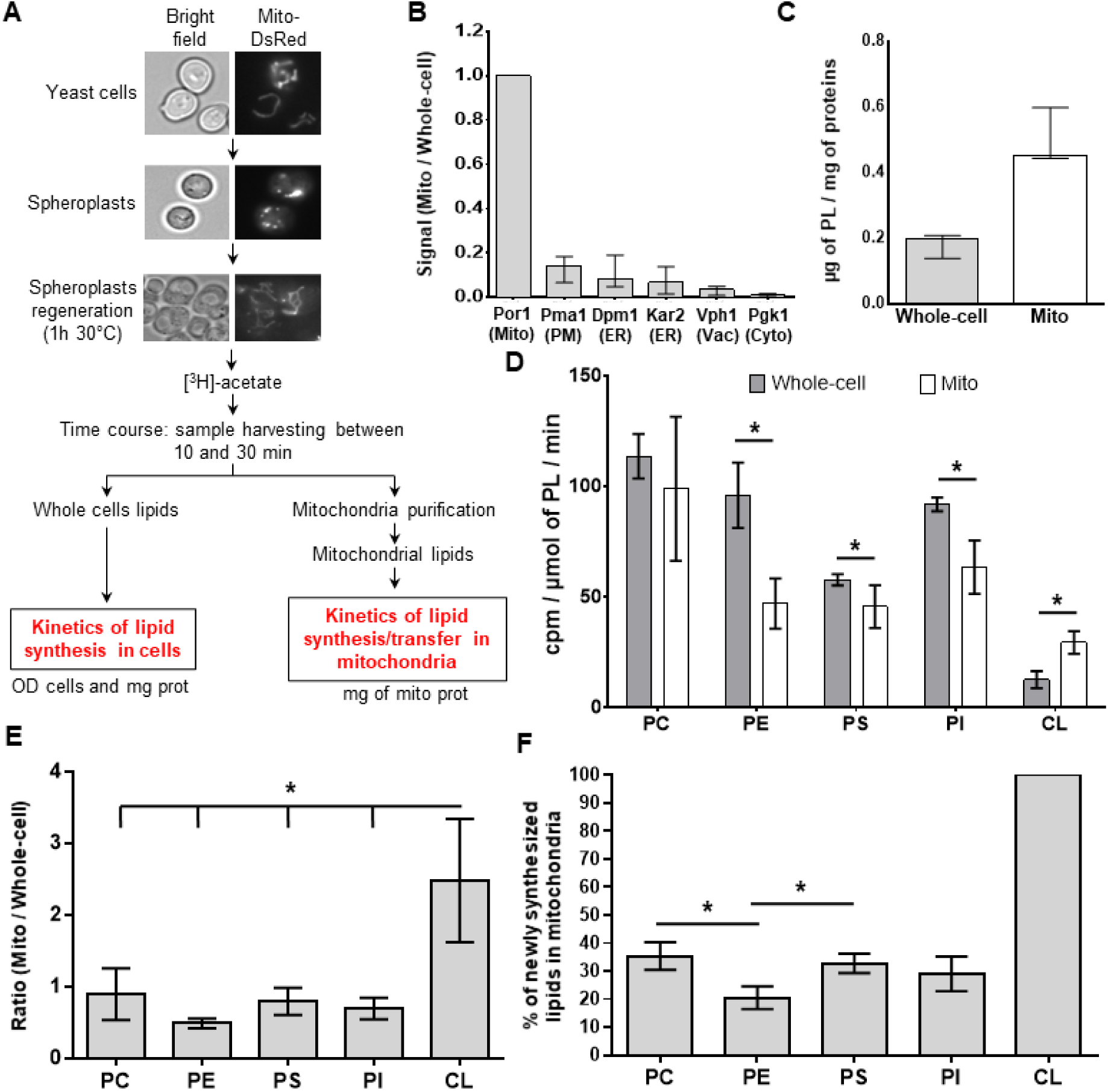
Quantification of *in vivo* lipid transport to mitochondria by metabolic radiolabeling. **A.** Schematic representation of the phospholipids-labeling assay. Yeast spheroplasts were allowed to recover for 1 h to restore mitochondrial morphology before labeling with [^3^H]-acetate. Samples were collected between 10 and 30 min after labeling to quantify radioactive phospholipids in whole-cell extracts and isolated mitochondria. **B.** Enrichment of mitochondrial proteins in isolated mitochondrial fractions. Immunoblot signals in mitochondrial fractions (Mito) were normalized to those measured in whole-cell. **C.** Phospholipids (PL) content in µmol normalized to protein abundance in mg in whole-cell and mitochondrial fractions. **D.** Rates of phospholipids labeling in whole-cell and mitochondrial fractions, expressed as counts per minute (cpm) per µmol of total phospholipids per minute. **E.** Ratio of phospholipids-labeling rates in mitochondrial fractions relative to whole-cell fractions. **F.** Estimated proportion of newly synthesized phospholipids present in mitochondria. Cardiolipin (CL), which is synthesized exclusively in mitochondria, was used as the mitochondrial reference (100%) to estimate the mitochondrial proportion of other phospholipid species. PC, phosphatidylcholine; PE, phosphatidylethanolamine; PI, phosphatidylinositol; PS, phosphatidylserine. Data are mean ± standard deviation from three independent biological replicates (n = 3). Statistical significance was determined using a Mann–Whitney test. *, p < 0.05.

The quality of mitochondrial preparations was assessed by immunoblotting. Mitochondrial fractions were strongly enriched in the mitochondrial marker Por1 relative to markers of the plasma membrane, ER, vacuole, and cytosol (**Fig. 1B**). For each sample, phospholipid content was normalized to protein abundance, allowing radioactivity incorporation to be expressed as counts per minute (cpm) per micromol of phospholipids and directly compared between whole-cell and mitochondrial fractions (**Fig. 1C**).

Using this assay, we monitored the incorporation of [^3^H]-acetate into phospholipid acyl chains in cells grown in rich medium. Radioactivity increased linearly over time for all major phospholipid classes except phosphatidic acid (PA), which displayed rapid turnover consistent with its role as a biosynthetic intermediate (**Fig. S1** and **S2**). Incorporation rates were determined from the slopes of individual labeling curves and expressed as cpm/µmol of phospholipids/min (**Fig. 1D**). As expected, cardiolipin (CL) exhibited a higher labeling rate in mitochondria than in whole-cell extracts, reflecting its exclusive synthesis within mitochondria. In contrast, labeling rates for phosphatidylserine (PS), phosphatidylethanolamine (PE), and phosphatidylinositol (PI) were lower in the mitochondrial fraction, whereas phosphatidylcholine (PC) showed similar rates in both fractions (**Fig. 1D**).

To estimate the proportion of newly synthesized phospholipids present in mitochondria, we first calculated the ratio of labeling rates in mitochondrial versus whole-cell fractions (**Fig. 1E**). Because CL is synthesized exclusively in mitochondria, we used it as an internal reference corresponding to 100% mitochondrial localization. This normalization enabled estimation of the mitochondrial fraction of each newly synthesized phospholipid classes (**Fig. 1F**). Approximately 30–35% of newly synthesized PC, PS, and PI molecules were detected in mitochondria, whereas the proportion was lower for PE (∼20%). This reduced value likely reflects the export of a substantial fraction of mitochondrially synthesized PE to other cellular membranes. Together, these data indicate that a substantial proportion of newly synthesized phospholipids transit through mitochondria shortly after their synthesis.

### In vivo measurement of lipid transport to mitochondria using heterologous galactolipid synthesis

Although the radiolabeling assay described above enables quantification of phospholipid transport to mitochondria, it does not readily address the substrate specificity of ER-mitochondria lipid transport pathways or allow estimation of lipid flux at the molecular level. To overcome these limitations, we developed an assay based on the synthesis of heterologous lipids that are absent from yeast cells. The principle of this approach is to synthesize a reporter lipid in the ER and monitor its transport to mitochondria through its conversion into a second lipid species by a mitochondrially localized enzyme. In this system, transport can be quantified directly from whole-cell lipid extracts without the need for mitochondrial purification. To ensure that transport is the rate-limiting step, the reporter lipids must be absent from yeast, the biosynthetic enzymes must be efficiently targeted to their respective compartments, and conversion of the transported lipid within mitochondria must occur rapidly relative to transport. Yeast cell membranes are mainly composed of phospholipids. Therefore, we selected the plant galactolipid pathway because galactoglycerolipids have not been detected in yeast. In plants, digalactosyldiacylglycerol (DGDG) is synthesized in two sequential reactions (Moellering and Benning, 2011; Boudière et al., 2014). First, monogalactosyldiacylglycerol (MGDG) is produced from diacylglycerol (DAG) and UDP-galactose by an MGDG synthase (MGD). MGDG is then converted into DGDG by a DGDG synthase (DGD) through the addition of a second galactose residue (**Fig. 2A**).

**Figure 2.**
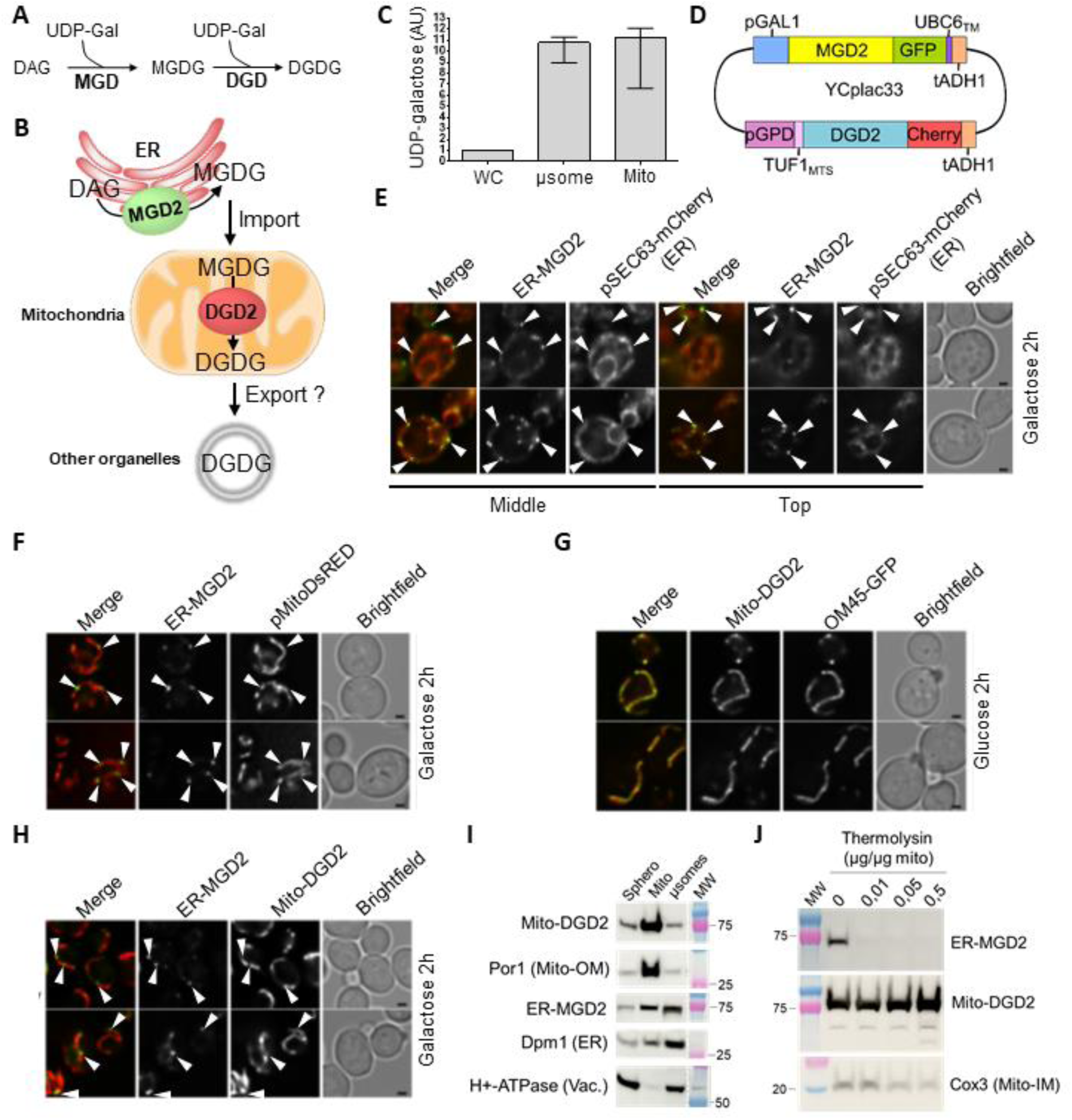
Development of a synthetic galactolipid-based assay to monitor ER to mitochondria lipid transport. **A.** Galactolipid synthesis pathway. Monogalactosyldiacylglycerol (MGDG) synthesis is mediated by a monogalactosyldiacylglycerol synthase (MGD) from diacylglycerol (DAG) and UDP-galactose (UDP-Gal). The subsequent addition of a galactose group on MGDG polar head allows the formation of digalactosyldiacylglycerol (DGDG) catalyzed by a digalactosyldiacylglycerol synthase (DGD). **B.** Schematic representation of the assay. *Arabidopsis thaliana* MGD2 was targeted to the ER to synthesize MGDG from DAG and UDP-galactose. DGD2 was targeted to mitochondria to convert ER-derived MGDG into DGDG. **C.** Relative abundance of UDP-galactose in whole-cell (WC), microsomal (µsome), and mitochondrial (Mito) fractions. UDP-galactose levels were normalized to whole-cell extracts. **D.** Schematic representation of the expression system. GFP-tagged MGD2 was anchored to the ER membrane through the transmembrane domain of Ubc6 (ER-MGD2). mCherry-tagged DGD2 was targeted to mitochondria using the N-terminal mitochondrial targeting sequence of Tuf1 (Mito-DGD2). ER-MGD2 expression was driven by the inducible *GAL1* promoter, whereas Mito-DGD2 was constitutively expressed from the *GPD1* promoter. Both expression cassettes were carried on the same YCplac33 plasmid. **E.** Colocalization of ER-MGD2 with the ER marker Sec63-mCherry following 2h of galactose induction. **F.** Localization of ER-MGD2 relative to the mitochondrial marker Mito-DsRed following 2h of galactose induction. **G.** Colocalization of Mito-DGD2 with the mitochondrial marker Om45-GFP in cells grown in glucose. **H.** Simultaneous visualization of ER-MGD2 and Mito-DGD2 after 2h of galactose induction. **I.** Subcellular fractionation analysis of ER-MGD2 and Mito-DGD2 localization. Spheroplast, mitochondrial (Mito), and microsomal (µsome) fractions were analyzed by immunoblotting using markers for mitochondria, the ER, and the vacuole. **J.** Protease protection assay of isolated mitochondria from ER-MGD2/Mito-DGD2 cells. Mitochondria were incubated with increasing concentrations of thermolysin, and protein sensitivity was assessed by immunoblotting. The mitochondrial inner membrane protein Cox3 served as a protected control. Data in B. represent mean ± standard deviation from three biological replicates.

We therefore used the well-characterized *Arabidopsis thaliana* enzymes MGD2 and DGD2 to establish a synthetic lipid transport assay in yeast. MGD2 and DGD2 are both membrane-bound enzymes (Kelly and Dörmann, 2002; Awai et al., 2001). As outlined in **Fig. 2B**, MGD2 was targeted to the cytoplasmic-facing surface of the ER membrane to drive MGDG synthesis and DGD2 was targeted to the mitochondrial matrix to convert MGDG that has been transported to the mitochondrial inner membrane to DGDG.

This strategy requires both that the enzymes are properly targeted and that the substrates the enzyme require are available in the ER and the mitochondrial matrix. MGD2 requires DAG, which is primarily synthesized in the ER (Klug and Daum, 2014), and UDP-galactose. Since MGDG also requires this substrate, we wanted to confirm it was available in both compartments. Metabolomic analysis of subcellular fractions revealed comparable enrichment of UDP-galactose in microsomal and mitochondrial fractions (**Fig. 2C**), indicating that substrate availability was unlikely to limit galactolipid synthesis.

After testing several targeting and expression strategies, we generated a system in which GFP-tagged MGD2 was anchored to the ER membrane through the transmembrane domain of Ubc6 (ER-MGD2), whereas mCherry-tagged DGD2 was directed to the mitochondrial matrix using the N-terminal targeting sequence of Tuf1 (Mito-DGD2) (**Fig. 2D**). Mito-DGD2 was expressed constitutively from the *GPD1* promoter (Bitter and Egan, 1984), whereas ER-MGD2 was expressed from the inducible *GAL1* promoter (Mumberg et al., 1994). Both expression cassettes were assembled on the same plasmid to ensure co-expression of the enzymes in all transformed cells (**Fig. 2D**). Fluorescence microscopy confirmed the expected localization of both enzymes. ER-MGD2 colocalized with the ER marker Sec63 and was distributed along the cortical and perinuclear ER (**Fig. 2E**). In addition, ER-MGD2 accumulated in discrete puncta that are frequently localized adjacent to mitochondria, suggesting the fusion protein may be enriched at ER-mitochondrial MCSs (**Fig. 2E and F**); the reason for this localization is not known. Mito-DGD2 showed complete overlap with the mitochondrial marker Om45, consistent with efficient mitochondrial targeting (**Fig. 2G**). Simultaneous visualization of both proteins confirmed their distinct localization patterns, although ER-MGD2 puncta were frequently observed in close proximity to Mito-DGD2-labeled mitochondria (**Fig. 2H**). Biochemical fractionation further supported these observations. Mito-DGD2 cofractionated with the mitochondrial marker Por1 and was strongly enriched in mitochondrial preparations (**Fig. 2I**). ER-MGD2 partitioned predominantly with microsomes and was also detected in mitochondrial fractions, but not more than the ER marker Dpm1. Protease protection assays demonstrated that Mito-DGD2 was protected from thermolysin digestion similarly to the mitochondrial inner membrane protein Cox3, whereas ER-MGD2 was readily degraded (**Fig. 2J**). These results confirm that ER-MGD2 remains exposed to the cytosol, whereas Mito-DGD2 is localized within mitochondria.

To determine whether the engineered galactolipid pathway was functional in yeast, we first analyzed galactolipid production following induction of ER-MGD2 expression. Thin-layer chromatography revealed the accumulation of MGDG in cells expressing ER-MGD2 and of DGDG in cells co-expressing ER-MGD2 and Mito-DGD2, whereas these lipids were not detected in control cells (**Fig. 3A**). These results demonstrate that both enzymes are active in yeast and that sufficient substrate is available in both compartments to support galactolipid synthesis. We next quantified galactolipid accumulation by lipidomics after 24 h of induction. MGDG and DGDG each reached levels above 40 mol% of total membrane glycerolipids, making them among the most abundant membrane lipids in induced cells (**Fig. 3B**). The high level of accumulation is consistent with the absence of known galactolipid catabolic pathways in yeast and indicates that galactolipid abundance primarily reflects biosynthesis and intracellular transport. To assess the physiological consequences of galactolipid production, we monitored cell growth under inducing and non-inducing conditions. No growth differences were observed in control media containing raffinose, which does not induce the *GAL1* promoter (**Fig. 3C**). Following galactose induction, all strains displayed similar growth kinetics during the first 10h. At later time points, MGDG production caused a modest reduction in growth rate, whereas DGDG production had a more pronounced effect (**Fig. 3C**). Together, these results establish that yeast cells efficiently synthesize and accumulate galactolipids and that high-level production, particularly of DGDG, affects cellular fitness.

**Figure 3.**
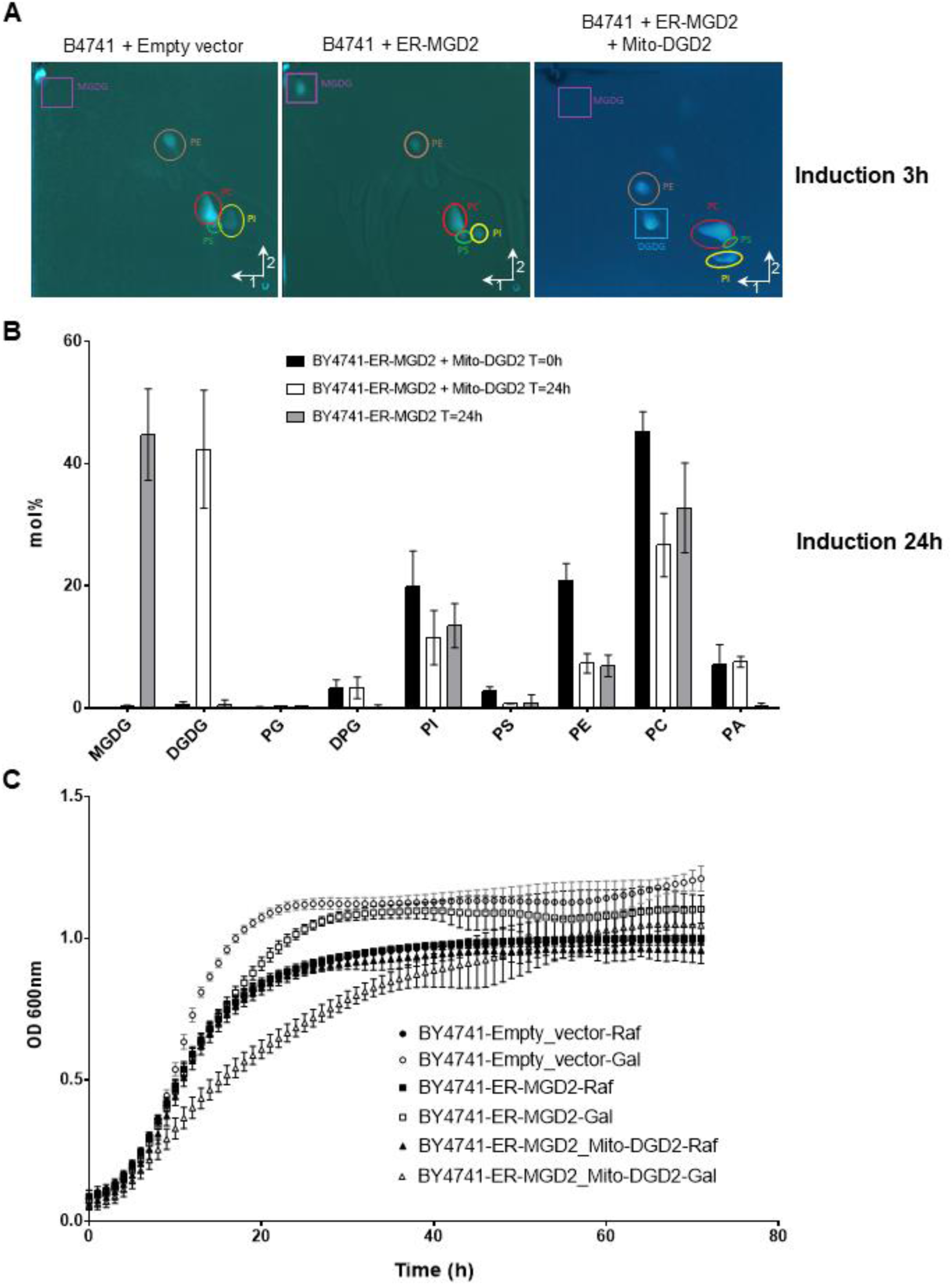
Functional analysis of the synthetic galactolipid pathway and its impact on yeast growth. **A.** Two-dimensional thin-layer chromatography analysis of lipids extracted from cells carrying an empty vector, ER-MGD2 alone, or ER-MGD2 together with Mito-DGD2 following 3h of galactose induction. Positions of monogalactosyldiacylglycerol (MGDG) and digalactosyldiacylglycerol (DGDG) are indicated by squares whereas phospholipids are indicated by circles. **B.** Lipidomic analysis of cells carrying an empty vector, ER-MGD2 alone, or ER-MGD2 together with Mito-DGD2 before induction (T = 0) and after 24 h of galactose induction. Glycerolipids classes abundance is expressed as mol% of total cellular glycerolipids. **C.** Growth analysis of cells carrying an empty vector, ER-MGD2 alone, or ER-MGD2 together with Mito-DGD2 under non-inducing (raffinose, Raf) and inducing (raffinose + galactose, Gal) conditions. CL, cardiolipin; PA, phosphatidic acid; PC, phosphatidylcholine; PE, phosphatidylethanolamine; PG, phosphatidylglycerol; PI, phosphatidylinositol; PS, phosphatidylserine. Data represent mean ± standard deviation from three independent biological replicates.

We next used this synthetic pathway to estimate lipid transport to mitochondria by monitoring the conversion of MGDG into DGDG. To ensure accurate measurement of transport rates, analyses were performed between 100 and 150 min after induction, a time window in which cell growth was unaffected and enzyme expression had reached a steady state (**Fig. 4A and B**). As expected, Mito-DGD2 remained constitutively expressed throughout the experiment, whereas ER-MGD2 expression stabilized approximately 110 min after galactose addition. Within this time window, DGDG accumulated linearly (R² = 0.94), increasing from approximately 3 to 8 mol% of total cellular lipids (**Fig. 4C**). In contrast, MGDG remained at very low abundance (<0.05 mol%) throughout the experiment. The near absence of MGDG despite continuous synthesis indicates its rapid conversion into DGDG and suggests that DGD2 activity is not the limiting step of DGDG production.

**Figure 4.**
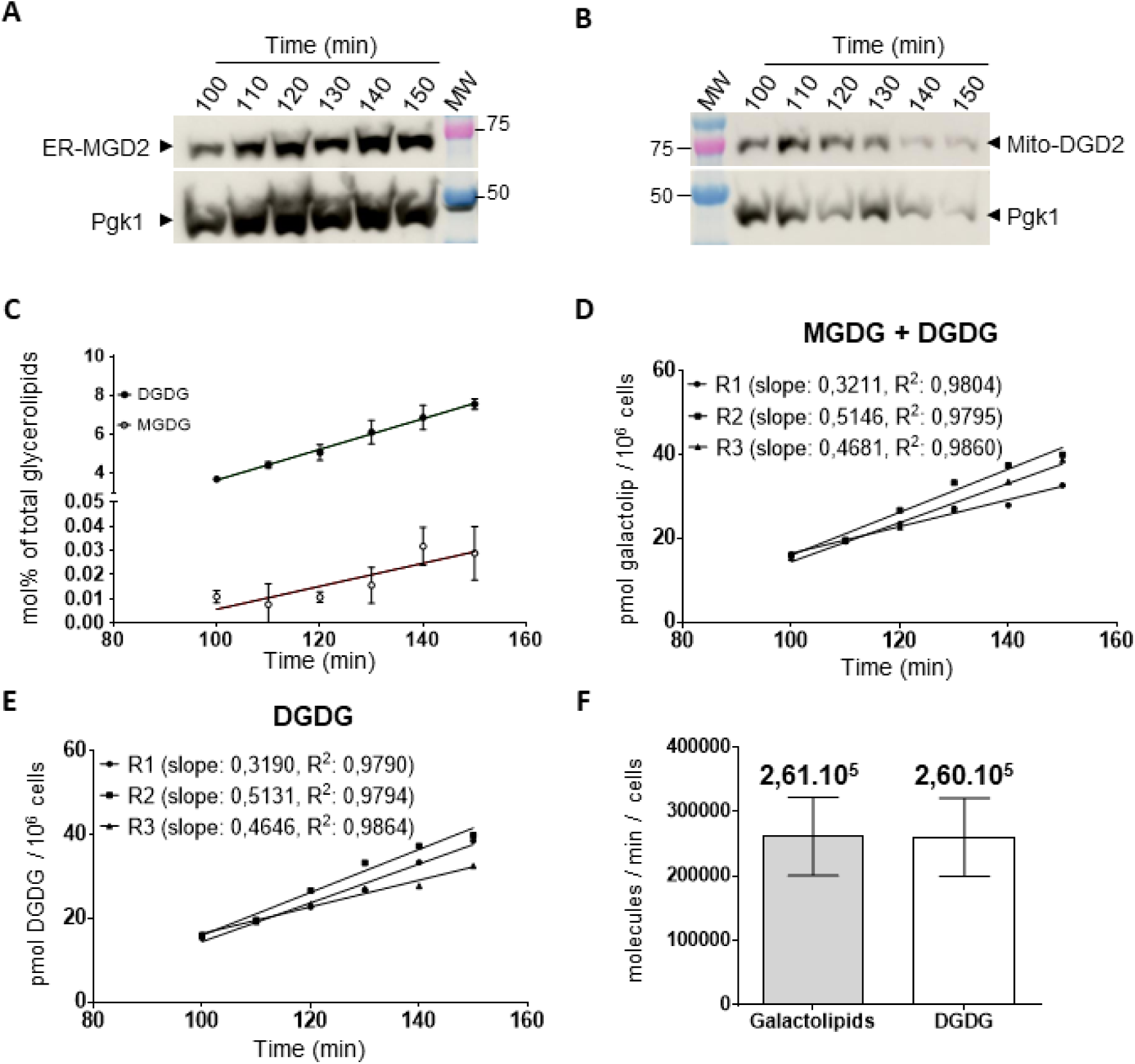
Estimated galactolipid synthesis and ER to mitochondrial lipid transport fluxes. A-B. Immunoblot analysis of ER-MGD2 (**A**) and Mito-DGD2 (**B**) expression during a 100–150 min time course following galactose induction. Pgk1 served as a loading control. **C.** Cellular abundance of monogalactosyldiacylglycerol (MGDG) and digalactosyldiacylglycerol (DGDG) during the time course, expressed as mol% of total cellular glycerolipids. **D.** Accumulation of total galactolipids (MGDG + DGDG) over time, expressed as pmol per 10^6^ cells. Linear regressions for the three independent biological replicates are shown together with the corresponding slopes and R² values. **E.** Accumulation of DGDG over time, expressed as pmol per 10^6^ cells. Linear regressions for the three independent biological replicates are shown together with the corresponding slopes and R² values. **F.** Average rates of total galactolipid synthesis and DGDG accumulation, expressed as molecules per cell per minute. Data in **C** and **F** represent mean ± standard deviation from three independent biological replicates (n = 3).

### To quantify transport flux, we calculated the accumulation rates of total galactolipids (MGDG

+ DGDG) and DGDG from three independent biological replicates (**Fig. 4D and E**). Total galactolipid synthesis occurred at a rate of 0.435 ± 0.101 pmol per 10^6^ cells per min, whereas DGDG accumulated at 0.432 ± 0.101 pmol per 10^6^ cells per min. Thus, >99% of newly synthesized MGDG was converted into DGDG during the measurement period. Conversion of these values into molecular fluxes yielded an estimated transport rate to mitochondria of 2.6 x 10^5^ MGDG molecules per cell per min (**Fig. 4F**). These findings indicate that lipid transfer from the ER to mitochondria can sustain remarkably high fluxes. Moreover, efficient transport of MGDG, a lipid absent from yeast throughout evolution, suggests that ER-mitochondria lipid transport pathways display limited substrate specificity and can accommodate multiple unrelated glycerolipids classes.

### Defining the rate and specificity of mitochondrial lipid transport in yeast

Using short-term metabolic labeling, we found that a substantial fraction of newly synthesized phospholipids rapidly appeared in mitochondrial fractions. Depending on the lipid class, approximately 20–35% of newly synthesized molecules were detected in mitochondria within minutes of synthesis. Although residual contamination of mitochondrial preparations by other cell compartments may contribute to a modest overestimation of these values, the results nonetheless indicate that lipid exchange between the ER and mitochondria occurs on a remarkably rapid timescale. These observations are consistent with the extensive physical coupling of both organelles through MCSs and support the view that mitochondria act as a major hub in cellular lipid metabolism (Mavuduru et al., 2024; Acoba et al., 2020; Tamura et al., 2020). Interestingly, the proportion of newly synthesized lipids detected in mitochondria differed among phospholipid classes (**Fig. 1F**). Whereas the proportion of newly synthesized PC, PS, and PI within mitochondria displayed similar ranges, the proportion of PE was significantly lower. One possible explanation is the rapid redistribution of mitochondrially-synthesized PE to other cellular membranes, where it contributes to PC biosynthesis and membrane homeostasis (Achleitner et al., 1995; Horvath and Daum, 2013).

To complement the radiolabeling approach, we established a synthetic transport assay based on the production of heterologous galactolipids. This system offers several advantages. First, transport can be quantified directly from whole-cell lipid extracts without requiring mitochondrial purification. Second, the directionality of transport is genetically defined by the localization of the biosynthetic enzymes, enabling selective monitoring of ER to mitochondria transfer, even if the indirect transport to mitochondria, as example through the vacuole, cannot be ruled out. Third, because galactolipids are absent from yeast, their abundance reflects *de novo* synthesis and transport without interference from endogenous metabolic pathways (*i. e.* degradation, modification of polar head…). Application of this assay revealed a transport flux of approximately 2.6 x 10^5^ lipid molecules per cell per minute. This value is remarkably high and is broadly consistent with previous estimates of mitochondrial lipid trafficking requirements during cell growth (*i. e.* 150 x 10^6^ molecules/cells/cell cycles) (Petrungaro and Kornmann, 2019). Together, these results provide a simple and quantitative framework for assessing lipid fluxes between organelles *in vivo*.

Newly synthesized MGDG accumulated only at trace levels, whereas DGDG production closely matched total galactolipid synthesis. These observations indicate that transport to mitochondria and subsequent conversion into DGDG occur with high efficiency and are unlikely to represent major rate-limiting steps under our experimental conditions. Instead, the overall flux measured by the assay appears to be primarily constrained by the rate at which MGDG is synthesized at the ER. This finding raises the possibility that mitochondrial lipid transport is closely coupled to lipid biosynthesis, providing a mechanism by which cells could coordinate growth, membrane production and inter-organelle lipid exchange.

A particularly notable finding is the efficient transport of galactolipids despite their absence from fungal membranes throughout evolution. MGDG headgroup composition differs substantially from endogenous yeast phospholipids, yet it was transferred to mitochondria with high efficiency. These observations suggest that some ER-mitochondria lipid transport pathways, likely mediated by bridge-like proteins, recognize membrane lipids with a highly limited substrate selectivity and might operate primarily through bulk membrane lipid extraction and transfer mechanisms rather than strict substrate-specific recognition.

Beyond the biological insights obtained here, the two assays described in this work provide complementary tools for the quantitative analysis of intracellular lipid transport in yeast. Whereas the radiolabeling assay enables monitoring of endogenous phospholipid trafficking, the synthetic galactolipid system permits direct measurement of transport flux and transport directionality. Because the subcellular localization of the biosynthetic enzymes can be readily modified, this strategy should be adaptable to the investigation of lipid exchange between a wide range of organelles. More broadly, these approaches provide a framework for dissecting the mechanisms, kinetics, and specificity of lipid transport pathways *in vivo*.

## Material and Methods

### Yeast strains and growth

The yeast strains used in this study are listed in **Table S1**. Yeast strains were grown in YPD media (yeast extract 1% (w/v), peptone 2% (w/v), glucose 2% (w/v)) or in selective CSM media (0,67% yeast nitrogen base with adenine (MP biomedicals), appropriate drop out media (MP biomedicals)) supplemented with glucose 2% (w/v) or raffinose 2% (w/v) for induction with galactose 2% (w/v).

### Cloning

All the cloning were performed using Gibson assembly according the protocol described in (Gibson et al., 2009). Briefly, overlapping PCR were performed and incubated with a linearized vector in a Gibson assembly mix before transformation in *E. coli*. PCR were performed with the Phusion DNA polymerase (Life technologies) in a final volume of 25 µL according to the manufacturer’s instructions. PCR products were separated on 1% agarose gels and bands were purified using the Monarch DNA gel extraction kit (New England Biolabs) according to manufacturer’s instructions. Vectors (500 ng) were digested in a final volume of 20 µL using the selected restriction enzymes (New England Biolabs).

Linearized vector and PCR products were mixed in the Gibson assembly mix and incubated 1h at 50°C. 5 µL were used to transform 25 µL of High efficiency NEB5α (New England Biolabs) competent cells according to the manufacturer’s instructions. Transformed plasmids were isolated using Miniprep kit (Qiagen) and plasmid sequences were checked by sanger sequencing.

### Yeast transformation

Yeast cells were transformed using a heat shock and lithium acetate protocol. Yeasts were grown overnight in the appropriate media and diluted 1/10 in the morning in the same media. After an incubation of 6h at 30°C under a 200 rpm agitation, yeasts were washed once in sterile water and once in LiAc-Sorbitol buffer (1M D-Sorbitol, 100 mM lithium acetate, 10 mM tris-acetate pH 8, 1 mM EDTA). Cells were resuspended in 250 µL of LiAc-Sorbitol buffer and 50 µL of cells were added to a mix containing 300 µL of LiAc-PEG (Polyethylene glycol 3350 40% (w/v), 100 mM Lithium acetate, 10 mM Tris-acetate pH 8, 1 mM EDTA), 5 µL of ssDNA from salmon sperm at 10 mg/mL and around 500 ng of plasmid. The mix was incubated 30 min at 30°C and 15 min at 42°C. Cells were centrifuged, resuspended in 200 µL of sterile water, spread on selective media and plates were incubated at 30°C. Transformed colonies were passed on a fresh plate and incubated at 30°C before doing experiments.

### In vivo radioactive labeling assay

Yeast cells were grown overnight at 30°C in 1L of YPD media until an OD600nm of 1 is reach. Cells were pelleted 10 min at 3 000 xg and resuspended in 50 mL of 100 mM Tris-S0_4_ pH 9,8, 10 mM DTT. Cells were incubated 10 min at 30°C under agitation at 100 rpm and pelleted 3 min at 3 000 xg. Cells were resuspended in 25 mL of **1.2M sorbitol buffer** (1.2 M Sorbitol, 20 % (v/v) YPD, 20 mM Hepes-KOH pH7.4) and Zymolyase® 20T (Amsbio) was added at a final concentration of 7.5 µg/OD. Spheroplasts were obtained by an incubation of 30 min at 30°C under agitation at 100 rpm. Spheroplasts were pelleted 3 min at 3 000 xg at 4°C and carefully washed twice with 5 mL of **1.2M sorbitol buffer**. For spheroplast regeneration, pellets were resuspended in 200 mL of YPD + 1.2 M sorbitol and incubated 1h at 30°C under an agitation of 100 rpm. Radioactive labeling was performed by the addition of 200 µL (1 µCi) of [H^3^]-Acetate (American Radiolabeled Chemicals ART 0202). At the different time points, 30 mL of spheroplasts (150 OD) were harvested and transferred into a pre-chilled Falcon containing 300 µL of 10 mM NaF and 300µL of 10 mM NaN_3_ to stop cell metabolism. One mL of the spheroplasts was harvested for total lipids extraction, a few µL for protein quantification by Bradford and the rest was used for mitochondria isolation. The pellets were resuspended in 1.5 mL of MIB (0.6 M sorbitol, 20 mM Hepes-KOH, Protease inhibitors tablet (Roche), 1 mM PMSF) and break by performing 50 strokes in a Potter. The pestle and potter were washed with 1.5 mL of MIB and the broken cells were centrifuged at 3 000 xg 5 min at 4°C. The supernatant was transferred into a clean tube and the pellet was resuspended in 1.5 mL of MIB and cells were broken again by 50 strokes in a Potter and washed with 1.5 mL of MIB. After a second centrifugation at 3 000 xg 5 min at 4°C, the supernatants were pooled and centrifuged at 10 000 xg 10 min at 4°C. The pellets were resuspended in 2 mL of MiB and centrifuged again at 3 000 xg 5 min at 4°C. The supernatants were centrifuged at 10 000 xg 10 min at 4°C and the pellets were resuspended in MIB and centrifuged again at 10 000 xg 10 min at 4°C. The pellets were resuspended in 2 mL of **SEM** (0,25 M sucrose, 10 mM MOPS-KOH pH7,2, 1 mM EDTA) and loaded on sucrose cushions (from bottom to top: 2 mL sucrose 60% (w/v), 4 mL sucrose 32% (w/v), 1,5 mL sucrose 23% (w/v) in 10 mM MOPS-KOH pH7,2, 1 mM EDTA). Tubes were centrifuged at 33 000 rpm 1h 4°C. Mitochondria ware harvested from the 60/32% interface and diluted in SEM buffer. Mitochondria were washed three time in SEM buffer by centrifugation of 10 min at 12 000 xg at 4°C. Mitochondria pellets were resuspended in SEM buffer. Proteins were quantified using BioRad Protein Reagent assay and lipids was then extracted and analyzed by 1D-TLC.

### Kinetic of galactolipid production in yeast

The yeast strain expressing ER-MGD2 + Mito-DGD2 was grown overnight at 30°C in 10 mL of CSM-Ura + raffinose media. At the end of the day, the preculture was used to start a 300 mL culture at a OD600/mL of 0.0035 in CSM-Ura + raffinose. Cells were grown overnight at 30°C 200 rpm until reaching an OD600 between 1 and 1.5. The experiment was started by the addition of galactose at 2% (w/v) in the media at T=0. At T=110, 120, 130, 140 and 150 min, 10 mL and 30 mL of cells were harvested for lipids and proteins extraction respectively. NaF 1 mM and NaN_3_ 1 mM were immediately added to stop the reactions and the OD600 was measured. Cells were then washed once in NaCl 0.9 % (w/v) and transferred in PreCellys tubes. Pellets were then immediately frozen in liquid nitrogen and stored at -70°C before processing.

### Lipid extraction

Yeast pellets were resuspended in 200 µL of sterile water and supplemented with 100 µL of acid-washed glass beads (425-500mm). Cells were broken at 4°C using a PreCellys Evolution cell lyser by 6 cycles of 20 sec at 10 000 rpm with a 30 sec break between each cycle. Broken cells were transferred into a hemolysis tube containing 3 mL of methanol/chloroform 2/1. Beads were washed with 600 µL of water and the supernatant was transferred in the methanol/chloroform 2/1 mix. For mitochondria lipid extraction, the mitochondria were resuspended in 800 µL of SEM and directly mixed with 3 mL of methanol/chloroform 2/1. The mix was vortexed 20 sec and incubated 10 min at RT before a centrifugation of 10 min at 1 500 rpm. The upper phase was discarded and 1 mL of KCl 2M was added to the lower phase. The mix was vortexed 20 sec and incubated 10 min at RT before a centrifugation of 10 min at 1 500 rpm. The lower phase was transferred in a clean hemolysis tube and 1 mL of chloroform was added to the upper phase. The mix was vortexed 20 sec and incubated 10 min at RT before a centrifugation of 10 min at 1 500 rpm. The lower phase was pooled with the first harvesting and lipids were dried under argon and stored at -20°C.

### Radioactive lipid analysis by 1D-TLC

Lipids were separated on 20x20 cm silica plates (Merck Millipore) in a solvent mixture chloroform/ethanol/water/triethylamine (30/35/7/35 (v/v)) according to the protocol described in (Vaden et al., 2005). After migration, TLC plates were dried and scanned on a RITA Star thin-layer analyzer (Raytest) to determine the count per minutes (cpm) for each phospholipid class at each time points. Using the µmol of total phospholipids / mg of proteins determined using the Bartlett assay, cpm/µmol of phospholipids was calculated at each time points. Slope were then calculated to obtain the velocity of incorporation of radioactivity in whole cells or mitochondria fraction in cpm/µmol of phospholipids/min (*Table S2*).

### Methanolysis and total fatty acid methyl esters (FAMEs) quantification by gas chromatography-flame ionization detector (GC-FID)

The lipid extracts are resuspended in 250 µL of ethanol-stabilized chloroform and 50 µL are mixed with 5 µg of internal standard (C15:0) and 3 mL of 2.5% sulfuric acid in methanol in sealable glass tubes and incubated at 100°C for 1 h to produce FAMEs. Reaction was stopped with 3 mL of water and phase separation was induced by the addition of 3 mL of hexane. After 20 min at room temperature, the upper organic phase containing the FAMEs was collected and the phase partition step is iterated. The collected hexane phases were evaporated under argon.

The FAMEs were resuspended in 100 µL of hexane and analyzed by GC-FID (Perkin Elmer Clarus 580) on a 30-m long cyanopropyl polysilphenesiloxane column (SGE BPX70) with a diameter of 0.22 mm and a film thickness of 0.25 μm. The GC column is heated at 180°C and nitrogen is used as vector gas. FAMEs were identified by comparison of their retention times with those of standards (Sigma-Aldrich) and quantified by the surface peak method using the C15:0 fatty acid internal standard for calibration.

### Lipid analysis by 2D-TLC

To analyze the presence in yeast expressing the different constructs, 150 µg of total lipids were loaded on a 10x10 cm silica HPTLC plate (Merck-Millipore). Samples were separated in the first dimension in the solvent mix A (chloroform 65 mL, methanol 25 mL, MilliQ water 4 mL). After drying the plate under a fume hood, the plate was turned at 90°C and lipids were separated in the second dimension in the solvent mix B (chloroform 50 mL, acetone 20 mL, methanol 10 mL, acetic acid 10 mL, MilliQ water 5 mL). After drying in the fume hood, the plates were spread with 8-Anilino-1-naphtalenesulfonic acid 0.2% (w/v) in methanol and lipids were revealed using a CAMAG TLC visualizer at 360 nm (CAMAG).

### Quantification of glycerolipids by high performance liquid chromatography coupled to triple quadrupole mass spectrometry (HPLC-MS/MS)

The lipid extracts corresponding to 25 nmol of FAMEs were dissolved in 100 µL of chloroform/methanol [2/1, (v/v)] containing 125 pmol of each internal standard. Internal standards used were PE 18:0-18:0 and DAG 18:0-22:6 (Avanti Polar Lipid) and SQDG 16:0-18:0 extracted from spinach thylakoid (Demé et al., 2014) and hydrogenated as described in (Buseman et al., 2006). Lipids were then separated by HPLC and quantified by MS/MS.

The HPLC separation method was adapted from (Rainteau et al., 2012). Lipid classes were separated using an Agilent 1260 Infinity II HPLC system using a 150 mm×3 mm (length × internal diameter) 5 µm diol column (Macherey-Nagel), at 40°C. The mobile phases consisted of hexane/isopropanol/water/ammonium acetate 1M, pH5.3 [625/350/24/1, (v/v/v/v)] (A) and isopropanol/water/ammonium acetate 1M, pH5.3 [850/149/1, (v/v/v)] (B). The injection volume was 20 µL. After 5 min, the percentage of B was increased linearly from 0% to 100% in 30 min and stayed at 100% for 15 min. This elution sequence was followed by a return to 100% A in 5 min and an equilibration for 20 min with 100% A before the next injection, leading to a total runtime of 70 min. The flow rate of the mobile phase was 200 µL/min. The distinct glycerophospholipid classes were eluted successively as a function of the polar head group.

Mass spectrometric analysis was done on a 6470 triple quadrupole mass spectrometer (Agilent) equipped with a Jet stream electrospray ion source under following settings: Drying gas heater: 260°C, Drying gas flow 13 L/min, Sheath gas heater: 300°C, Sheath gas flow: 11L/min, Nebulizer pressure: 25 psi, Capillary voltage: ± 5000 V, Nozzle voltage ± 1000. Nitrogen was used as collision gas. The quadrupoles Q1 and Q3 were operated at widest and unit resolution respectively. PC analysis were carried out in positive ion mode by scanning for precursors of m/z 184 at a collision energy (CE) of 35eV. PE, PI, PS, PG, PA, MGDG and DGDG measurements were performed in positive ion mode by scanning for neutral losses of 141 Da, 277 Da, 185 Da, 189 Da, 115 Da, 179 Da and 341 Da at CEs of 29 eV, 21 eV, 21 eV, 25 eV, 25 eV, 8 eV and 11 eV, respectively. Quantification was done by multiple reaction monitoring (MRM) with 30 ms dwell time. DAG and TAG species were identified and quantified by MRM as singly charged ions [M+NH_4_]+ at a CE of 19 and 26 eV respectively with 30 ms dwell time. CL species were quantified by MRM as singly charged ions [M-H]- at a CE of -45 eV with 50 ms dwell time. Mass spectra were processed by MassHunter Workstation software (Agilent) for identification and quantification of lipids. Lipid amounts (pmol) were corrected for response differences between internal standards and endogenous lipids and by comparison with a quality control (QC). QC extract corresponds to a known lipid extract from yeast cells containing galactolipids qualified and quantified by TLC and GC-FID as described by (Jouhet et al., 2017).

### Phospholipids quantification using the Bartlett assay

Phospholipids were quantified from total or mitochondrial lipid extracts of known protein quantity. Lipids were resuspended in chloroform and 20 µL and 30 µL of lipids were transferred in a glass tubes 16x125 mm and dried under nitrogen. 125 µL of Acid solution (60 mg vanadium IV-oxide, 40 mL sulfuric acid, 20 mL perchloric acid) were added to the samples and the tubes were heated on gas flame until the color becomes orange and a white smoke is seen. After cooling down the mix, 2.5 mL of Ammonium molybdate solution (0.1 g ammonium molybdate, 0.313 g Powder Mix (15 g sodium metabisulfite, 0.5 g sodium sulfite, 0.25 g 4-amino-3-hydroxy-1-naphtalensulfonic acid) in 50 mL H_2_O were added and samples were heated at 100°C for 8 min. After cooling down the mix, the OD at 800 nm was measured and the nmol of phosphate per mg of cells or mitochondria was calculated according to a standard curve performed with a 1 mM K_2_HPO_4_ solution.

### Growth curves

Yeast growth analyses were performed on TECAN Spark. Yeasts were grown overnight at 30°C in CSM-Ura + raffinose at 30°C under 200 rpm. Cells were diluted at a OD600 nm of 0.03 in a final volume of 200 µL of CSM-Ura + raffinose (non-induced) or CSM-Ura + galactose (induced). Cultures were transferred into a 96-wells transparent plate. The growth was followed by measuring the OD600 nm every 15 min after a shaking of 15 sec at 100 rpm. Three technical replicates were performed per plates and the experiment was carried out in three biological replicates.

### Protein extraction

For protein extraction, yeast pellets (around 30 OD) were resuspended in Extraction buffer (8M Urea, 50 mM Tris-HCl pH6.8, 1 mM EGTA, 1 mM DTT). Cells were broken at 4°C using a PreCellys Evolution cell lyser by 3 cycles of 20 sec at 10 000 rpm with a 30 sec break between each cycle. Broken cells were centrifuged 5 min at 5 000xg at 4°C and the supernatant was transferred in a clean tube. Protein quantification was performed using Bradford Protein Assay (BioRad).

### Western Blots

Twenty µg of proteins were mixed with Bolt^TM^ LDS Sample buffer 4X and Bolt^TM^ Sample reducing agent 10X reducing reagent (Life technologies), heated at 95°C for 5 min and loaded of Bolt^TM^ 4-12% Bis-tris pre-cast gels (Life technologies). Migration was performed at 120V in MES SDS Bolt^TM^ migration buffer (Life technologies) and transfer was performed in 0.22 µm nitrocellulose membrane 1h at 90V in transfer buffer (193 mM glycine, 25 mM Tris-HCl pH 8.3, ethanol 20% (v/v)). Membrane was stained with Ponceau Red (ponceau red 0.2 % (w/v), trichloroacetic acid 3 % (v/v)) and destained in TBS-T (Tris-HCl 50 mM pH 7.5, NaCl 150 mM, Tween 20 0,1% (v/v)). Western blots were performed in TBS-T plus milk 5% (w/v) and revealed using the Clarity or Clarity Max (BioRad). The antibodies used are the following: GFP-HRP (Miltenyi Biotech), mCherry (Biolegend), Por1, Dpm1, Pgk1, Vph1, Kar2, Pma1 (Invitrogen).

### Subcellular fractionation

To analyze the subcellular localization of proteins, a yeast pre-culture in CSM-URA + raffinose was cultured overnight at 30°C with a 200rpm agitation. Three 2L flask of 500 mL of CSM-URA + raffinose were then prepared at OD600=0.02 and grown overnight at 30°C with a 200rpm agitation. The transgene expression was induced by the addition of galactose 2% (w/v) final and performed during 1h at 30°C under a 200rpm agitation. OD cells was measured and spheroplast formation, cells breaking and mitochondria isolation was performed as described in the “*In vivo radioactive labeling assay*” part with adaptation of the buffers’ volumes according to total OD600 cells and with slight modification: the 1.2M sorbitol buffer was modified to continue induction (1.2 M Sorbitol, 20% (v/v) CSM-URA + galactose 2% (w/v), 20 mM Hepes-KOH pH7.4) and spheroplasting was performed during 70 min. To isolate the microsomal fraction, the supernatant from the first 10 000 xg centrifugation step was harvested and centrifuged 1h at 100 000 xg at 4°C. The pellet was resuspended in SEM buffer and used for protein quantification using the BioRad Protein Assay Reagent (BioRad).

### Thermolysin treatments

For thermolysin treatment, isolated mitochondria were washed twice in SM buffer (sucrose 0.25M, 10mM MOPS-KOH pH7.2). The reaction was performed in a final volume of 50 µL with 10 µg of mitochondria supplemented with 0, 0.1, 0.5 or 5µg of thermolysin from *Geobacillus stearothermophilus* (Sigma Aldrich) in SM buffer supplemented with 1 mM CaCl_2_. Samples were incubated on ice for 1h and the reaction was stopped by the addition of Bolt^TM^ LDS Sample buffer 4X and Bolt^TM^ Sample reducing agent 10X reducing reagent (Life technologies) and heating 5 min at 95°C. Samples were then analyzed by western blot.

### Fluorescence imaging

Cells were imaged live on a Nikon brightfield Ti2E motorized inverted microscope (Nikon), with a triggered Piezo Z PINano P-736 for Z stage control (Physik Instrumente LP), spectra III 8 line LED light source (Lumencor), CFI60 plan apochromat lambda D 100x oil immersion objective lens (Nikon), Oko black enclosure set at 30°C (Okolab), and Kinetix 22 back-illuminated sCMOS (Teledyne Photometrics) camera. The system was operated with NIS-Elements software (Nikon).

### Metabolomic analysis of UDP-galactose

To determine UDP-galactose levels in mitochondrial, cytosol, and microsome fractions of *Saccharomyces cerevisiae*, a standard metabolite extraction protocol was applied. In this procedure, 1 mL of cold acetonitrile/methanol/water (4:4:2, v/v/v) was added to each sample. To improve metabolite recovery, samples were vortexed, sonicated for 5 min, and incubated at −20°C for 1 h. Extracts were evaporated to dryness using a Thermo Scientific Savant SpeedVac concentrator and reconstituted in 50 µL of Milli-Q® water prior to ion chromatography–high-resolution mass spectrometry (IC-HRMS) analysis. The analysis was performed using an IC-HRMS platform consisting of a Dionex™ ICS-5000+ Reagent-Free™ HPIC™ system (Thermo Fisher Scientific, Sunnyvale, CA, USA) coupled to a Q Exactive Plus Orbitrap mass spectrometer (Thermo Fisher Scientific, Waltham, MA, USA) equipped with a heated electrospray ionisation (HESI) source. Anion-exchange chromatography was carried out using an IonPac AS11-HC analytical column (250 × 2 mm) coupled to an AG11-HC guard column (50 × 2 mm) (Dionex). The system was equipped with an eluent generator (ICS-5000+ EG) allowing on-line generation of potassium hydroxide (KOH). Separation was achieved at a flow rate of 0.38 mL/min using a linear KOH gradient over a total run time of 50 min. The gradient program was as follows: 7 mM KOH for 1.0 min; linear increase from 7 to 15 mM over 8.5 min; hold for 10.5 min; increase to 45 mM over 10 min; increase to 70 mM over 3 min; increase to 100 mM over 0.1 min; hold for 8.9 min; decrease to 7 mM over 0.5 min; and re-equilibration at 7 mM for 7.5 min. The column temperature was maintained at 25°C, and the autosampler temperature at 6°C. The injection volume was 15 µL. High-resolution mass spectrometry was performed in negative ionisation mode using full-scan Fourier transform mass spectrometry (FTMS) at a resolution of 70,000 (at m/z 200). The HESI source parameters were as follows: capillary temperature, 380°C; auxiliary gas heater temperature, 380°C; sheath gas flow rate, 50 arbitrary units; auxiliary gas flow rate, 10 arbitrary units; S-Lens RF level, 80%; and spray voltage, 2.75 kV. UDP-galactose was identified and quantified by exact-mass extraction with a mass tolerance of 5 ppm. All measurements were performed in triplicate using independent biological samples.

## Supplemental material

**Figure S1:**
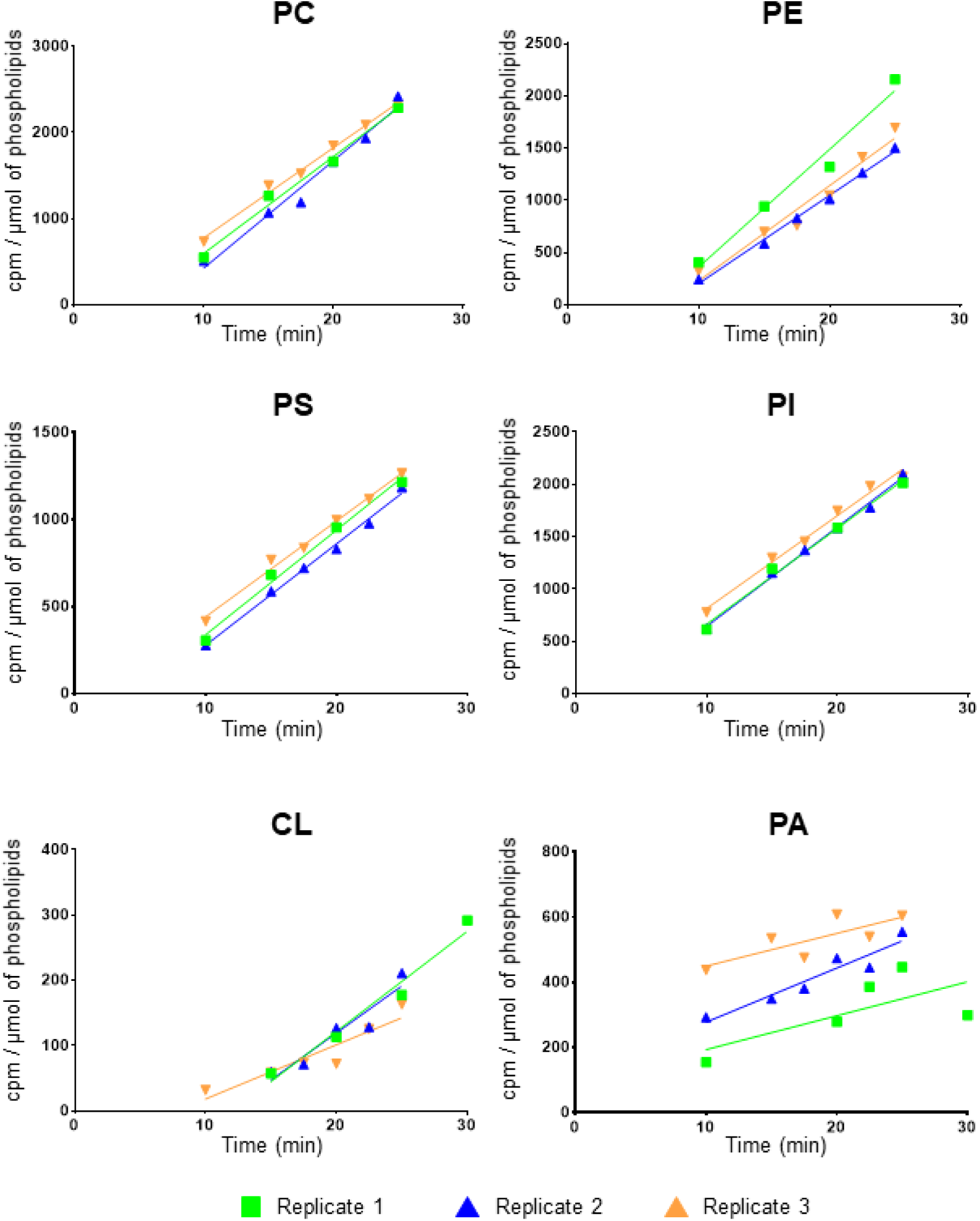
Incorporation of radioactivity (in count per min (cpm)) into phospholipids of the whole-cell extract over time in the three biological replicates performed.

**Figure S2:**
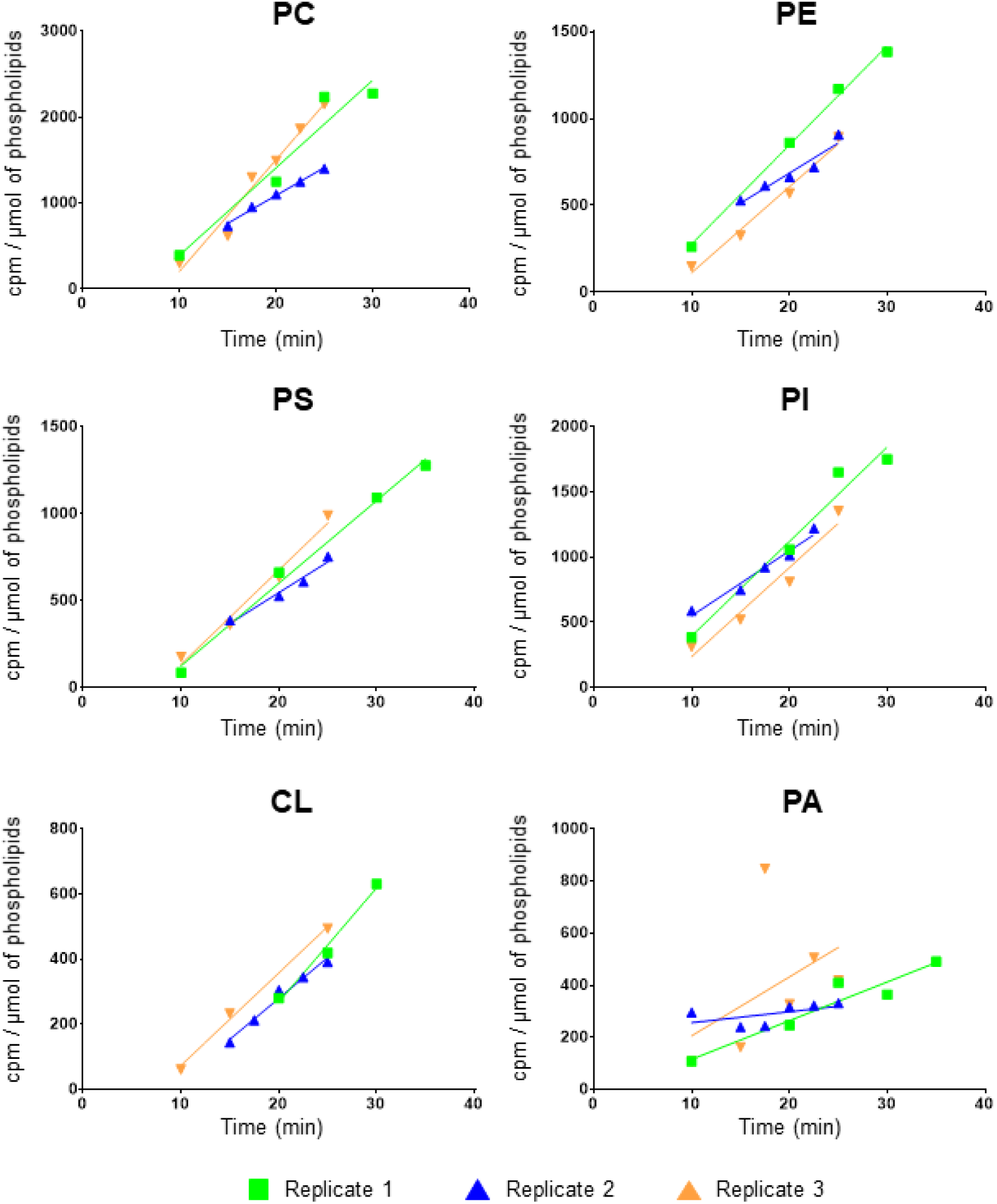
Incorporation of radioactivity (in count per min (cpm)) in phospholipids of mitochondrial extract over time in the three biological replicates performed.

**Table S1:**
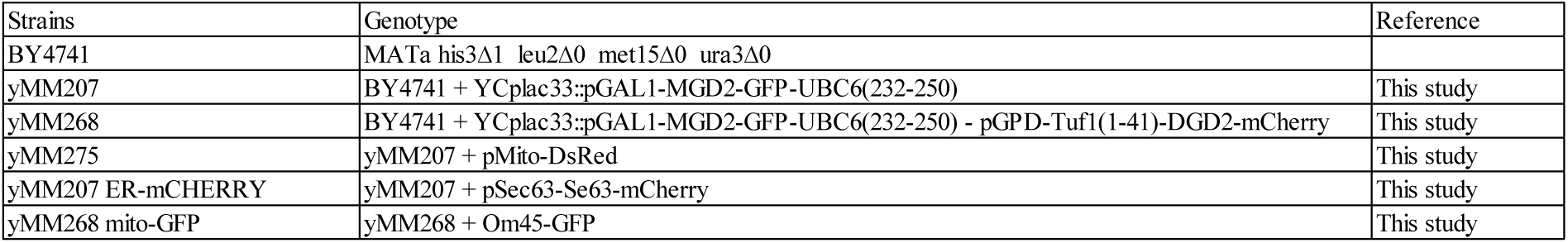
List of yeast strains used in this study.

| Strains | Genotype | Reference |
| --- | --- | --- |
| BY4741 | MATa his3Δ1 leu2Δ0 met15Δ0 ura3Δ0 |  |
| yMM207 | BY4741 + YCplac33::pGAL1-MGD2-GFP-UBC6(232-250) | This study |
| yMM268 | BY4741 + YCplac33::pGAL1-MGD2-GFP-UBC6(232-250) - pGPD-Tuf1(1-41)-DGD2-mCherry | This study |
| yMM275 | yMM207 + pMito-DsRed | This study |
| yMM207 ER-mCHERRY | yMM207 + pSec63-Se63-mCherry | This study |
| yMM268 mito-GFP | yMM268 + Om45-GFP | This study |

**Table S2:**
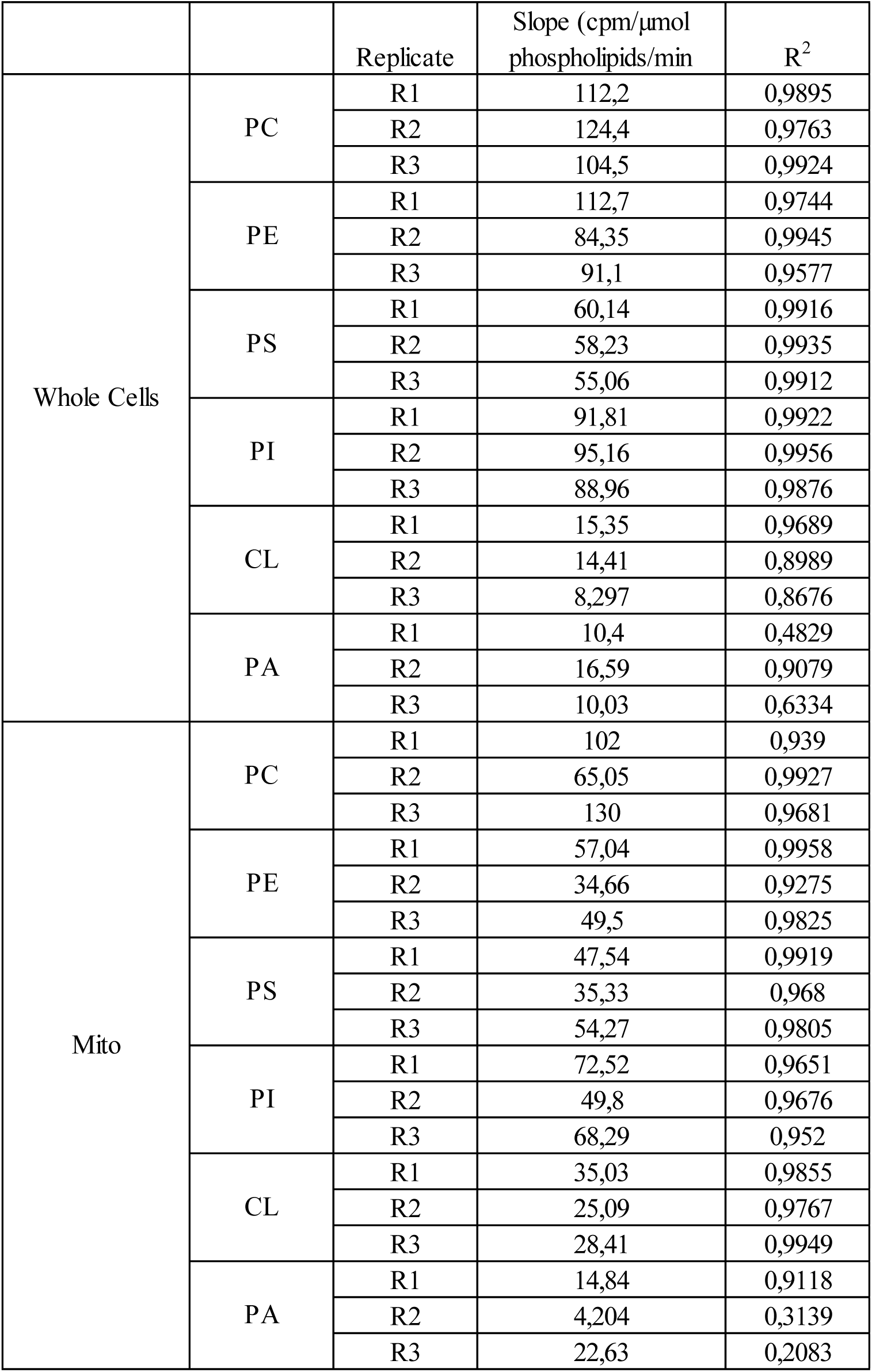
Slopes and R^2^ obtained for each replicate and each lipid classes in whole-cell and mitochondrial fractions.

| | | Replicate | Slope (cpm/μmol<br>phospholipids/min | R<sup>2</sup> |
| --- | --- | --- | --- | --- |
| Whole Cells | PC | R1 | 112,2 | 0,9895 |
|  |  | R2 | 124,4 | 0,9763 |
|  |  | R3 | 104,5 | 0,9924 |
|  | PE | R1 | 112,7 | 0,9744 |
|  |  | R2 | 84,35 | 0,9945 |
|  |  | R3 | 91,1 | 0,9577 |
|  | PS | R1 | 60,14 | 0,9916 |
|  |  | R2 | 58,23 | 0,9935 |
|  |  | R3 | 55,06 | 0,9912 |
|  | PI | R1 | 91,81 | 0,9922 |
|  |  | R2 | 95,16 | 0,9956 |
|  |  | R3 | 88,96 | 0,9876 |
|  | CL | R1 | 15,35 | 0,9689 |
|  |  | R2 | 14,41 | 0,8989 |
|  |  | R3 | 8,297 | 0,8676 |
|  | PA | R1 | 10,4 | 0,4829 |
|  |  | R2 | 16,59 | 0,9079 |
|  |  | R3 | 10,03 | 0,6334 |
| Mito | PC | R1 | 102 | 0,939 |
|  |  | R2 | 65,05 | 0,9927 |
|  |  | R3 | 130 | 0,9681 |
|  | PE | R1 | 57,04 | 0,9958 |
|  |  | R2 | 34,66 | 0,9275 |
|  |  | R3 | 49,5 | 0,9825 |
|  | PS | R1 | 47,54 | 0,9919 |
|  |  | R2 | 35,33 | 0,968 |
|  |  | R3 | 54,27 | 0,9805 |
|  | PI | R1 | 72,52 | 0,9651 |
|  |  | R2 | 49,8 | 0,9676 |
|  |  | R3 | 68,29 | 0,952 |
|  | CL | R1 | 35,03 | 0,9855 |
|  |  | R2 | 25,09 | 0,9767 |
|  |  | R3 | 28,41 | 0,9949 |
|  | PA | R1 | 14,84 | 0,9118 |
|  |  | R2 | 4,204 | 0,3139 |
|  |  | R3 | 22,63 | 0,2083 |

## Acknowledgment

We thank Jodie Nunnari for providing pMito-DsRED plasmid. This work was supported by the French National Research Agency in the framework of the “investissement d’avenir” program (ANR-15-IDEX-02), NIH no. R35GM153315 (to W.A.P.), and the Intramural Research Program of the National Institute of Diabetes and Digestive and Kidney Diseases (no. 1 ZIA DK060004 to W.A.P.). The LIPANG (Lipid analysis in Grenoble) platform is supported by the Rhône-Alpes Region, the fonds FEDER, and GRAL, financed within the University Grenoble Alpes graduate school (Ecoles Universitaires de Recherche) CBH-EUR-GS (ANR-17-EURE-0003). We thank the MetaboHUB-MetaToul facility (ANR-11-INBS-0010) for metabolomic analyses of UDP-galactose content.

